# Audiovisual detection at different intensities and delays

**DOI:** 10.1101/173773

**Authors:** Chandramouli Chandrasekaran, Matthias Gondan

## Abstract

In divided-attention tasks with two classes of target stimuli (e.g., auditory and visual), redundancy gains are typically observed if both targets are presented simultaneously, as compared with single-target presentation. Different models explain such redundant-signals effects, including race and coactivation models. Here we generalize one such coactivation model, the superposition model, and show that restricting this evidence accumulation process to a limited time interval, that is a temporal deadline, provides the model the ability to describe the detection behavior of two monkeys across a wide range of intensities and stimulus onset asynchronies. We present closed-form solutions for the mean absorption times and probabilities for the two-stage diffusion process with drift towards a single barrier in the presence of a temporal deadline.

## Background

Integrating information from different senses improves perception and action, speeds up response time (RT), enhances detection and discrimination accuracy, and facilitates correct arm movements (Cluff, Crevecoeur, & Scott, 2015; Crevecoeur, Munoz, & Scott, 2016; Diederich & Colonius, 2004a; Miller, 1982; Nickerson, 1973; Sakata, Yamamori, & Sakurai, 2004; Schwarz, 1989, 1994; Seilheimer, Rosenberg, & Angelaki, 2014; Stein & Stanford, 2008; van Atteveldt, Murray, Thut, & Schroeder, 2014). Despite many studies of multisensory integration, several questions about how the behavioral benefits of multisensory integration emerge remain unanswered (Chandrasekaran, 2016; Cluff et al., 2015; Crevecoeur et al., 2016).

In this study, we focused on the question of how the behavioral benefits of multisensory integration vary as a function of both the sensory intensities as well as delays between the stimuli. With few exceptions (Gondan, Götze, & Greenlee, 2010), past studies have largely investigated these two aspects in isolation (Cluff et al., 2015; Crevecoeur et al., 2016; Diederich & Colonius, 2004a; Dixon & Spitz, 1980; Holmes, 2009; Meredith, Nemitz, & Stein, 1987; Stein, Meredith, Huneycutt, & McDade, 1989; van Wassenhove, Grant, & Poeppel, 2007). In this study, we extended our recent paradigm where monkeys detect visual, auditory, and audiovisual vocalizations in a background of noise (Chandrasekaran, Lemus, & Ghazanfar, 2013; Chandrasekaran, Lemus, Trubanova, Gondan, & Ghazanfar, 2011; Miller, 1986) by incorporating delays between the sensory stimuli (Miller, 1986). We measured the response times (RTs) and detection accuracy of the monkeys and then modeled the behavioral patterns by expanding on the coactivation modeling framework (Diederich, 1995; Miller, 1982; Schwarz, 1989, 1994, 2006).

We demonstrate that the behavioral benefits accrued by monkeys when detecting vocalizations are a lawful function of both the intensities and the delay between sensory stimuli (Cluff et al., 2015; Crevecoeur et al., 2016; Dixon & Spitz, 1980; Holmes, 2009; Meredith et al., 1987; Stein et al., 1989; van Wassenhove et al., 2007). After ensuring that separate activation models are insufficient to explain the behavior of the monkeys (Miller, 1986; Raab, 1962) we show that the superposition framework that assumes additivity of visual and auditory inputs and accumulation to a bound provides an approximate description of the RTs but fails to describe the accuracy of the monkeys performing the task (Diederich, 1995; Schwarz, 1989, 1994). We, therefore, expanded on the diffusion superposition model by incorporating a deadline for accumulation. We derive closed-form solutions for the mean reaction times (RTs) and accuracy rates for the two-stage diffusion process with drift towards a single barrier in the presence of a temporal deadline. We then show that such an expansion of the classical coactivation model provides an excellent description of both accuracies as well as RTs of these monkeys.

The manuscript is organized as follows. We provide a brief background on the diffusion superposition model. We then describe the details of and results from our behavioral experiments in monkeys and show that the classical model provides an unsatisfactory fit to the accuracy and response times. We then show that extending the model by incorporating a deadline for accumulation provides an better account of the observed behavior of the monkeys. We end by discussing the ramifications of this model.

### Redundant Signals Tasks

The redundant signals task is a simple and powerful experimental paradigm for investigating multisensory perception. In this task, subjects are asked to respond in the same way to stimuli of two sensory modalities, for example, auditory and visual stimuli (A, V, Diederich & Colonius, 1987; Miller, 1982, 1986). The typical finding is that if signals from both modalities are present (redundant signals, AV), average responses are faster than average responses to targets from any single modality (unimodal targets). This redundant signals effect, by itself, is not necessarily indicative of any special mechanism that integrates the information of the different channels. Redundancy gains may arise from a race between the channel-specific detection processes (“race model,“Miller, 1982; Raab, 1962). In audiovisual detection, however, redundancy gains are typically larger than predicted by race models and these are thought to be better explained by “coactivation models” that assume some kind of integration of the information provided by the two sensory systems (Diederich, 1995; Diederich & Colonius, 2004b; Miller, 1986; Schwarz, 1989, 1994). One such model is the diffusion superposition model, which we explain below.

### The diffusion superposition model

The single-barrier diffusion superposition model (Schwarz, 1994) is a coactivation model that describes redundancy gains assuming additive superposition of channel-specific diffusion processes. In diffusion models, the assumption is that the presentation of a stimulus leads to a buildup of evidence that is described by a noisy diffusion process **X**(*t*) with drift *μ* and variance σ^2^ > 0 (Ratcliff, 1988; Ratcliff & McKoon, 2008; Ratcliff, Smith, Brown, & McKoon, 2016; Smith & Ratcliff, 2004). The stimulus is ‘detected’ as soon as an evidence criterion *c*> 0 is met for the first time. The density *g(c)* and distribution *G(t)* of the first-passage times **D** are well known (‘inverse Gaussian’, Cox & Miller, 1965),

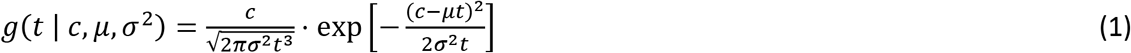

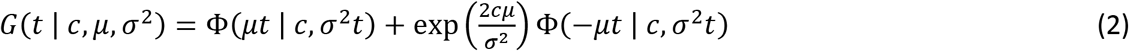

with Φ(*x|m,s^2^*) denoting the Normal distribution with mean *m* and variance *s^2^* detection time *E*(**D**) is obtained by integrating 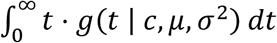 which simplifies to

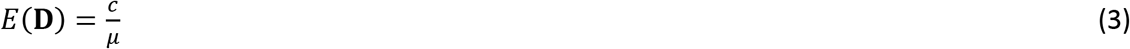

Predictions for the detection times for unimodal stimuli A and V are, therefore, easily obtained, *c/μ_A_* and *c/μ*_V_, respectively. When two stimuli are presented simultaneously, coactivation occurs. The model assumes that the two modality-specific processes superimpose linearly, **X**_AV_(*t*) =**X**_A_(*t*) +**X**_V_(*t*). The new process **X**_AV_ is, therefore, again a diffusion process with drift *μ_AV_* = *μ*_A_ + *μ*_V_ and variance 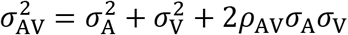 (under the assumption that **X**_A_ and **X**_V_ are uncorrelated, the covariance term will be zero). Since the drift parameters add up, **X**_AV_ reaches the response criterion earlier than any of its single constituents, resulting in faster responses to redundant stimuli,

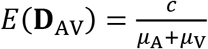

What happens in stimuli presented with onset asynchrony (SOA, e.g., V160A, i.e., the visual stimulus component is presented first, and then the auditory stimulus component follows the visual stimulus component with a delay τ, here 160 ms)? During the first τms, the sensory evidence is accumulated by the visual channel alone. If the criterion is reached within this interval, the stimulus is detected, and a response is initiated. This case occurs with probability

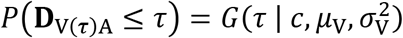 given by Eq. (2). If detection occurs before a time of has elapsed, it is expected to occur within

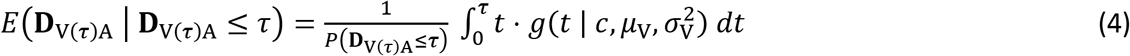

The solution for the integral is given by Schwarz (1994, Eq. 6). In the other case, the process has attained a subthreshold activation level **X**(τ) =*x* <*c*, with probability density described by

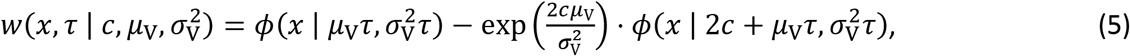

(e.g., Schwarz, 1994, Eq. 7) with *ϕ*(x|μ,σ^2^) denoting the Normal density with mean μ and variance σ^2^. Because activation level *x* has already been attained, less ‘work’ has to be done to reach the criterion. Starting at time τ, both stimuli contribute to the buildup of activity, resulting in an aggregate process drifting with μ_AV_ towards a residual barrier *c*-*x*:

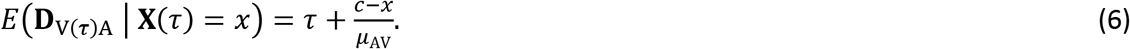

This expectation must be integrated for all possible levels of activation *x* < *c*, weighted by the density (5) that level *x* has been reached by time *t* = τ:

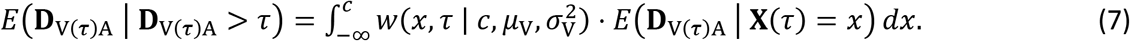

An analytic solution for the overall, unconditional expected first-passage time *E*(**D**_V(τ)A_) has been derived by Schwarz (1994, Eq. 10).

The diffusion process only describes the ‘detection’ latency **D**, or processing time. To derive a prediction for the observed response time **T**, an additional variable **M** is typically introduced that summarizes everything not described by the diffusion process (e.g., motor execution), such that **T**=**D** + **M**. Therefore, the model prediction for the mean response time is

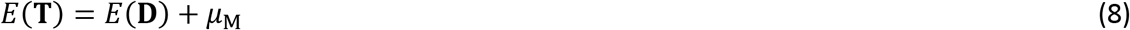

where the additional parameter denotes the expectation of **M**. Schwarz (1994) demonstrated that such a model of additive superposition quite accurately describes the mean response times and variances reported for Participant B.D. from (Miller, 1986) in a speeded response task with 13 different SOAs (at a single intensity).

### Present behavioral experiment

Our stated goal is to understand how the benefits in redundant signals tasks depend on both the intensity of the sensory stimuli as well as the SOA. To that end, we trained monkeys to detect visual, auditory and audiovisual vocal signals in a constant background of auditory noise (Chandrasekaran et al., 2013; Chandrasekaran et al., 2011). We chose a free response paradigm (without explicit trial markers, Shub & Richards, 2009) because it mimics natural audiovisual communication—faces are usually continuously visible and move during vocal production. The task was a typical redundant signals task (Miller, 1982, 1986); stimuli were chosen to approximate natural face-to-face vocal communication. This task was heavily inspired by the observation that besides providing benefits for discrimination of speech sounds (Besle, Fort, Delpuech, & Giard, 2004; Grant, Walden, & Seitz, 1998), visual speech enhances the detection of auditory speech (Grant, 2001; Grant & Seitz, 2000; Schwartz, Berthommier, & Savariaux, 2004). In such settings, the vocal components of the communication signals are degraded by environmental noise. The motion of the face, on the other hand, is usually perceived clearly. In the task, monkeys detected the onset of ‘coo’ calls that are affiliative vocalizations commonly produced by macaque monkeys in a variety of contexts (Hauser & Marler, 1993; Rowell & Hinde, 1962). The coo calls were presented at three different levels of sound intensity and were embedded in a constant background noise. The visual signals were videos of monkey avatars opening their mouth to make a coo vocalization. The size of the mouth opening was in accordance with the intensity of the associated vocalization: greater sound intensity was coupled to larger mouth openings by the dynamic face. Finally, we generated audiovisual stimuli by combining the videos with the coo vocalizations. The task of the monkeys was to detect the visual motion of the mouth or the onset of the coo vocalization. Audiovisual stimuli were presented either in synchrony or at 10 different SOAs.

## Methods

### Subjects

Nonhuman primate subjects were two adult male macaques (S and B, *Macaca fascicularis*). The monkeys were born in captivity and housed socially. The monkeys underwent sterile surgery for the implantation of a painless head restraint (see Chandrasekaran, Turesson, Brown, & Ghazanfar, 2010). All experiments and surgical procedures were performed in compliance with the guidelines of the Princeton University Institutional Animal Care and Use Committee.

### Procedure

Experiments were conducted in a sound attenuating radio frequency enclosure. The monkey sat in a primate chair fixed 74 cm opposite a 19 inch CRT color monitor with a 1280 × 1024 screen resolution and 75 Hz refresh rate. The screen subtended a visual angle of ~25° horizontally and 20° vertically. All stimuli were centrally located on the screen and occupied a total area (including blank regions) of 640 × 653 pixels. For every session, the monkeys were placed in a restraint chair and head-posted. A depressible lever (ENV-610M, Med Associates) was located at the center-front of the chair. Both monkeys spontaneously used their left hand for responses. Stimulus presentation and data collection were performed using Presentation (Neurobehavioral Systems).

### Stimuli

Coo vocalizations could be one of three different levels of sound intensity (85 dB, 68 dB, 53 dB) and were embedded in a constant background noise of ~63 dB SPL (giving us a range of signal to noise ratios, SNR, Fig. 1A). We used coo calls from two macaques as the auditory components of vocalizations; these were recorded from individuals that were unknown to the monkey subjects. The auditory vocalizations were resized to a constant duration of 400 ms using a MATLAB implementation of a phase vocoder (Flanagan & Golden, 1966) and normalized in amplitude (Fig. 1A). The visual components of the vocalizations were 400 ms long videos of synthetic monkey agents making a coo vocalization. The animated stimuli were generated using 3D Studio Max 8 (Autodesk) and Poser Pro (Smith Micro), and were extensively modified from a stock model made available by DAZ Productions (Silver key 3D monkey, Figs. 1B, C). Further details of the generation of these visual avatars are available in a prior study (Chandrasekaran et al., 2011).

**Figure 1.**
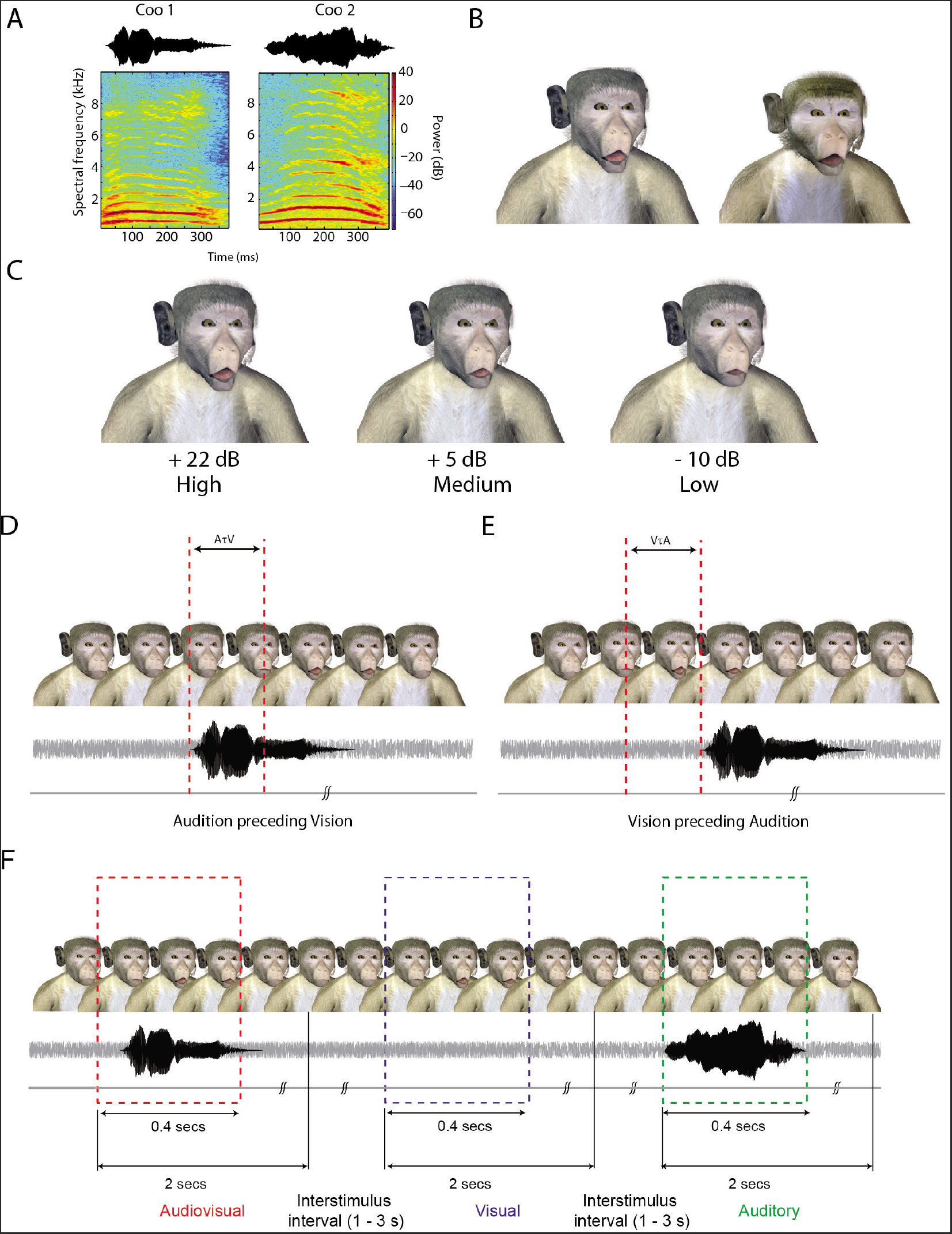
Stimuli, task structure, and behavior. **A:** Waveform and spectrogram of coo vocalizations detected by the monkeys. **B:** Frames of the two monkey avatars at the point of maximal mouth opening for the largest SNR. **C:** Frames with maximal mouth opening from one of the monkey avatars for three different SNRs of +22 dB (High), +5 dB (Medium) and –10 dB (Low). **D:** An illustration of the A(*τ*)V audiovisual condition. In this stimulus, the onset of the vocalization precedes the onset of the mouth motion. **E:** An illustration of the V(*τ*)A audiovisual condition. In this stimulus, visual cues such as mouth opening precede the onset of the vocalization. **F:** Task structure for monkeys. An avatar face was always on the screen. Visual, auditory and audiovisual stimuli were randomly presented with an inter stimulus interval of 1–3 seconds drawn from a shifted and truncated exponential distribution. Responses within a 2 sec window after stimulus onset were considered hits. Responses in the inter stimulus interval are considered to be response errors and led to timeouts.

The audiovisual stimuli were generated by presenting both the visual and auditory components either in synchrony or at 10 different stimulus onset asynchronies (SOAs). Audition could precede vision by 240, 160, 120, 80 or 40 ms (Fig. 1D) and vice versa (Fig. 1E). Intensities were always paired; that is the weak auditory stimulus was always paired with a small mouth opening. We, therefore, had 3 pairs of intensities and 11 SOAs, which resulted in 33 AV conditions in total. Each block also had 3 auditory intensities, 3 visual intensities. Catch trials (C) were used to discourage from spontaneous responses and to be able to control for fast guesses in the analysis of the RT distributions.

### Task

During the task (Fig. 1F), an avatar face (e.g., Avatar 1) was continuously present on the screen; the background noise was also continuous. In the visual-only condition (V), Avatar 1 moved its mouth without any corresponding auditory component. In the auditory-only condition (A), the vocalization paired with Avatar 2 was presented with the static face of Avatar 1. Finally, in the audiovisual condition (AV), Avatar 1 moved its mouth accompanied by the corresponding vocalization of Avatar 1 and in accordance with its intensity. We, therefore, had two AV stimuli. (A1V1 and A2V2). In the even blocks, the avatar face was the still frame of V1, A2 was the auditory sound played, and A1V1 was the audiovisual stimulus. The other block had the opposite configuration. This task design avoids the conflict between hearing a vocalization with the corresponding avatar face not moving.

Stimuli of each condition (V, A, AV, C) were presented after a variable inter stimulus interval between 1 and 3 seconds (drawn from a shifted and truncated exponential distribution). Monkeys indicated the detection of a V, A or AV event by pressing the lever within 2 seconds following the onset of the stimulus. In the case of hits, the ISI was started immediately following a juice reward. In the case of misses, the ISI began after the two second response window.

After every block of 126 trials (33 AV stimuli × 3 trials + 3 A × 3 trials, 3 V × 3 trials, 9 catch trials), a brief pause (~10 to 12 seconds) was imposed. Then, a new block was started in which, the avatar face, and the identity of the coo sound used for the auditory-only condition were switched. Within a block, all the conditions were randomly interleaved with one another.

### Training

Monkeys were initially trained over many sessions to respond to the coo vocalization events in visual, auditory or audiovisual conditions while withholding responses when no stimuli were presented. A press of the lever within a window starting 150 ms after onset of the vocalization event and within two seconds led to a juice reward and was defined as a hit. An omitted response in this two-second window was classified as a miss similar to the studies of free response tasks (without explicit trial markers, Shub & Richards, 2009). Lever presses to catch trials were defined as false alarms. In addition, the random presses during the interstimulus interval (ISI) were discouraged by enforcing a timeout where no stimuli were presented. The timeout was chosen randomly from a distribution between 3.0 and 5.5 s. The monkeys had to wait for the entire duration of this timeout period before a new stimulus was presented. Any lever press during the timeout period led to a renewal of the timeout with the duration again randomly drawn from the same distribution. Monkeys were trained until the erroneous presses in this ISI period were less than or equal to 10% of trials in any given session.

### Statistical analysis of behavioral performance

Hit rate was defined as the ratio of hits to the total number of targets. For each SNR and condition, the accuracy was defined as the ratio of hits to hits plus misses expressed as a percentage. The false alarm rate (i.e., presses to the catch stimuli) was defined as the number of false alarms divided by the total number of catch trials. Mean RTs and SDs were computed for the correct responses; confidence intervals were obtained by resampling the observed RT distributions (including omitted responses and false alarms) with replacement 1000 times and estimating the standard deviation of the mean of the resampled data.

### Test of the race model inequality

An important model class for redundant signals effects is the so-called race model, or separate activation model (Colonius & Diederich, 2006; Gondan & Minakata, 2016; Miller, 1982, 1986; Raab, 1962). According to the race model, redundancy benefits are not due to an actual integration of visual and auditory signals but because of parallel processing of both signals. In the bimodal stimulus, the two channels engage in a race-like manner (“parallel first-terminating model”, Townsend & Ashby, 1983), so that the probability for fast responses is increased because slow processing times are canceled out by the other channel. The redundancy gains obtained in the race model are limited, however, and a classical test of whether this mechanism can explain the observed reaction times is the well-known race model inequality (Miller, 1982), stating that the RT distribution for redundant stimuli *F*_AV_(*t*) never exceeds the sum of the RT distributions for the unisensory stimuli *F*_A_ (*t*), *F*_V_ (*t*),

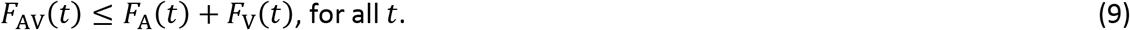

(Eriksen, 1988, see also Gondan and Heckel, 2008) demonstrated that this inequality could be refined by taking into account anticipatory responses to catch trials (C),

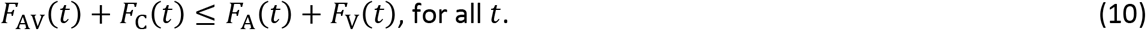

For redundant targets presented with SOAτ, the inequality generalizes to (Miller, 1986)

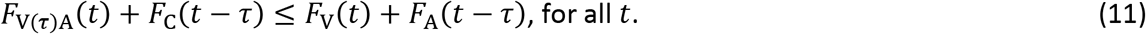

If this inequality is violated in a given data set, then parallel self-terminating processing cannot account for the benefits observed for multisensory stimuli, suggesting an explanation based on the integration of the signals (e.g., the superposition model described above; see also (Luce, 1986; Miller, 2016) for a discussion of the assumption of context independence).

For each condition, we determined the empirical cumulative distribution functions (eCDFs) and then computed the maximum violation, that is, the maximum difference between the left-hand side and the right-hand side of Inequality 11. A bootstrap technique (Miller, 1986), was used to assess the statistical significance of the observed violations of the race model inequality.

### Test of the diffusion superposition model

We fit the predictions of the diffusion superposition model to the mean RTs from the two monkeys. We used the analytic equation provided in Schwarz (1994, Eq. 10). To perform model fitting, we computed for each monkey, an approximate 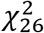 goodness-of-fit statistic given by the sum of the squared standardized deviations of the predicted and the observed average response times (e.g., Schwarz, 2006). The 26 degrees of freedom are given by the difference between the number of conditions (3 intensities × 13 SOAs) minus the number of model parameters: 12 parameters are due to the drift and variance for each visual and auditory SNR (3 SNRs × 2 modalities × 2 parameters). The thirteenth parameter is the average residual non-decision time (Eq. 8).

## Results

Figures 2A, C show the accuracy of the two monkeys in the detection task for different SNRs and SOAs. The accuracy (see Figure 2A, C, see also Supplementary Tables S1–S4) showed lawful decreases with changes in SNR for the auditory components of vocalizations (e.g., only about 55% detection rate with auditory stimuli). In contrast, changes in mouth opening size for the visual component of the vocalizations, which were meant to match the changes in auditory intensity, had only minimal impact on the accuracy of the animals. The mean RTs (Figures 2B,D) showed the wing shaped pattern typically observed in redundant signals tasks with SOA manipulations (Miller, 1986; Ulrich & Miller, 1997).

**Figure 2:**
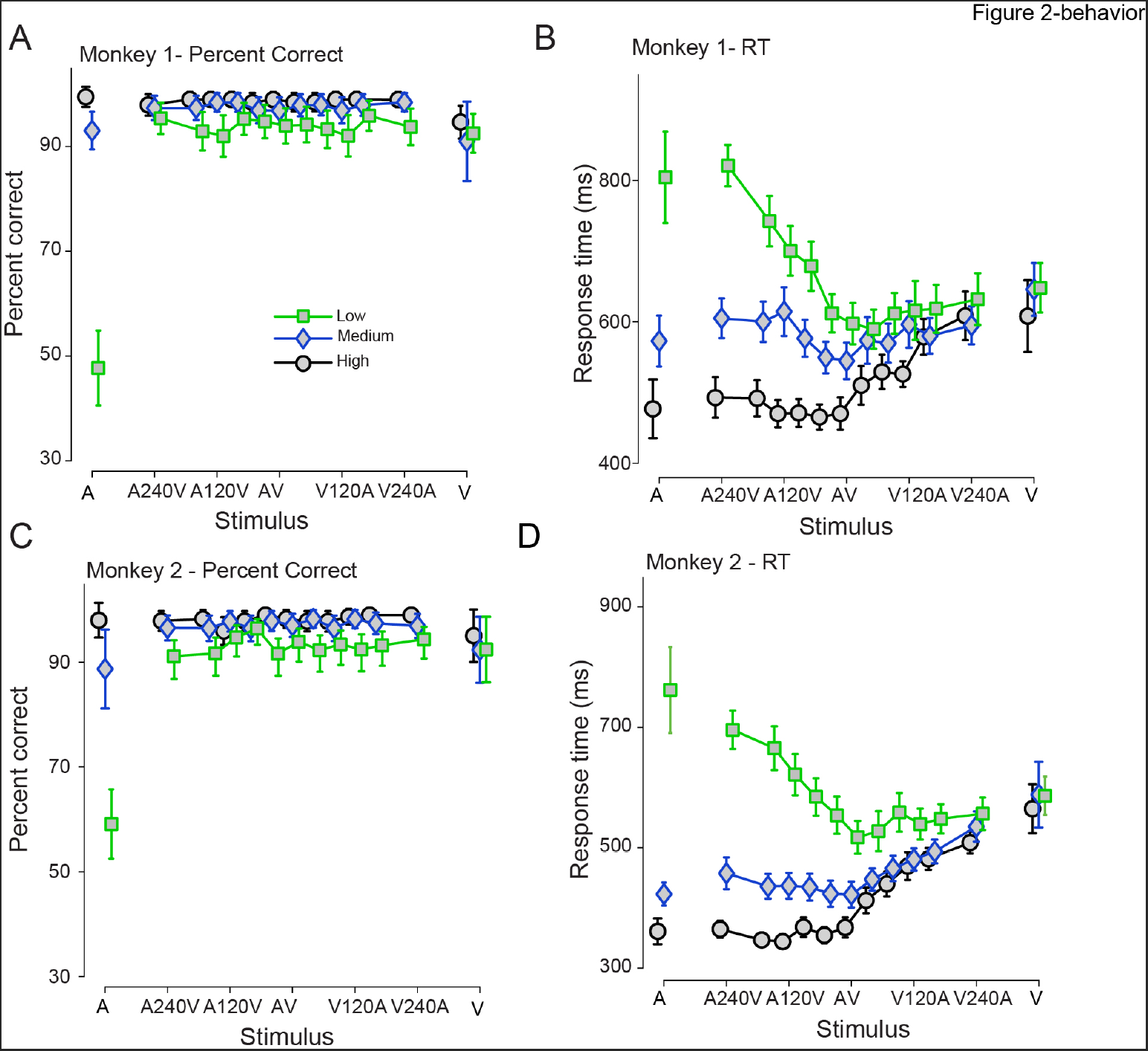
Monkeys integrate visual and auditory cues across a large range of SOAs and SNRs. A, B: Mean accuracy and RTs for audiovisual, visual-only and auditory-only conditions for Monkey 1 (A, B) and Monkey 2 (C, D) as a function of different stimulus conditions and SNR. X-axes depict stimuli. Y-axes in the left panel depict the accuracy in percent. Y-axes in the right panel denote response times in ms. Error bars denote confidence intervals around the mean (2 × SEM).

In both monkeys, statistically significant violations of the upper bound of the race model inequality for RTs (Inequality 11) occurred for a large range of SOAs, indicating that the observed redundancy gains were inconsistent with a race model. For Monkey 1, statistically significant violations of the RMI (at the criterion of *p* < 0.05) were observed in 28 out of the 33 multisensory conditions, for Monkey 2, this was observed in all 33 conditions (Tables S5, S6 in online supplement).

We first examined if the diffusion superposition model could describe the behavior of the monkeys during this task. We found that the superposition model could describe the overall pattern of the mean RTs of the monkeys (Figures 3A, B). However, the overall fit was unsatisfactory and especially poor for the lowest SNRs (Monkey 1: 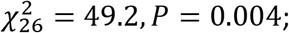 Monkey 2: 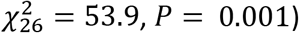

**Figure 3:**
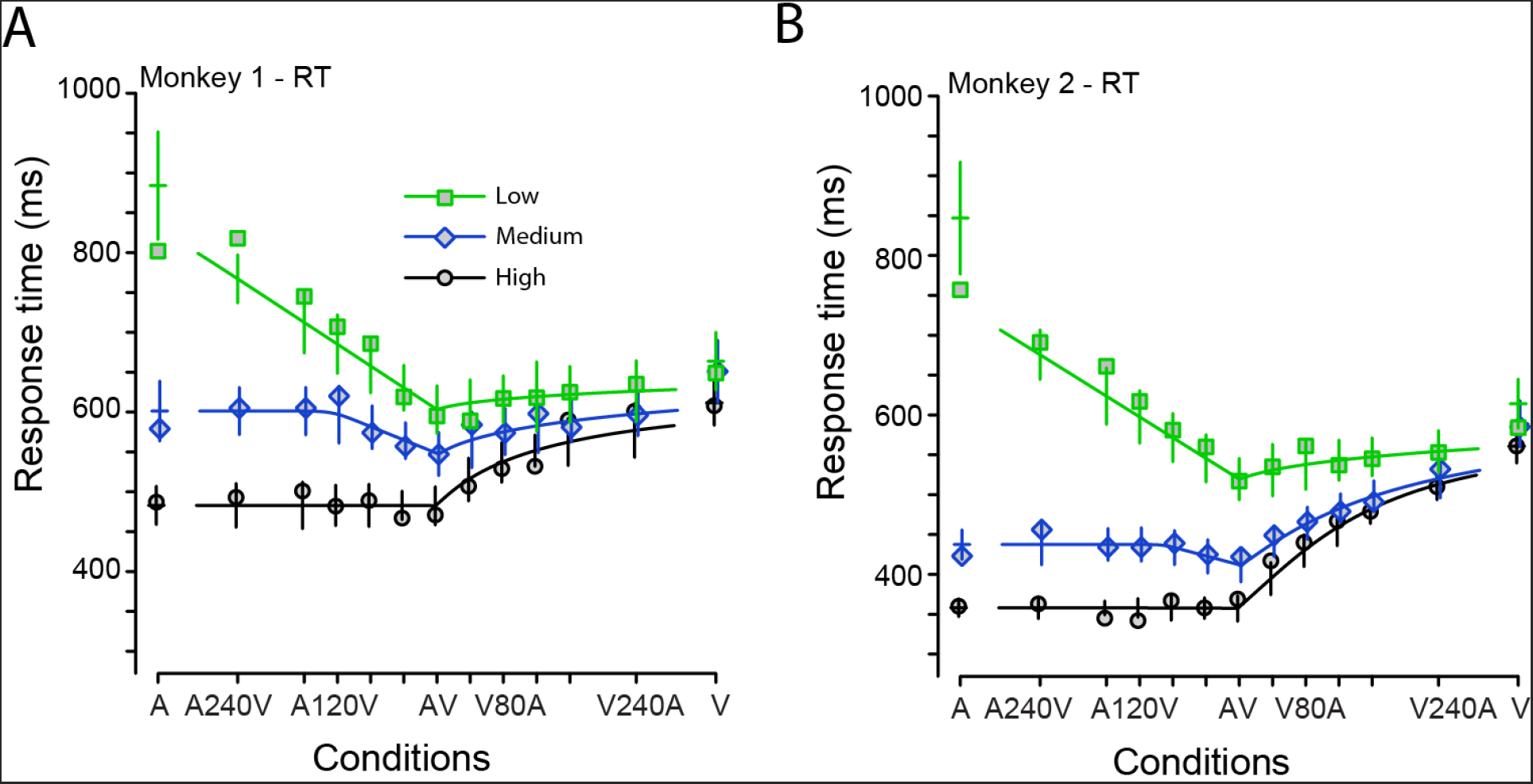
The diffusion superposition model provides an incomplete description of the detection behavior of monkeys. Audiovisual, visual-only and auditory-only RTs for both monkeys along with the predicted RT shown in lines according to the diffusion superposition model as a function of SNR (squares = low, diamonds = medium, circles = high)and SOA. Error bars denote confidence intervals (2 × standard error) around predicted mean RTs. A shows the RT for monkey 1; B shows the RT for monkey 2.

The monkeys also omitted a substantial proportion of responses to weak auditory stimuli (detection accuracy rates for the lowest auditory intensity were ~55% and 60%), which is, by design, not accounted for by the superposition model described above. The integral in Equation 3 ranges from 0 to infinity, such that, absorption at the upper barrier is a certain event given enough time. This means that the superposition model always predicts ceiling level accuracies for all intensities, a prediction clearly inconsistent with the observed behavioral data.

### A coactivation model with a deadline

An unrealistic assumption of the model described above is that accumulation will always complete, which in the single-barrier diffusion model implies that the monkeys have 100 percent detection accuracy. Given enough time, a diffusion process with drift μ > 0 will almost certainly reach the criterion. From an experimental perspective, this has several implications: the intensity of the stimulus components must be sufficiently high to ensure detection rates of 100% and the temporal window for responding is infinitely long to guarantee that all responses are collected. However, if the temporal window for stimulus detection is limited by a deadline (we assume that *d* > τ) the proportion of correct responses is given by the distribution of the detection times at *t*=*d*. For unimodal stimuli and synchronous audiovisual stimuli, this probability corresponds to the inverse Gaussian distribution at time 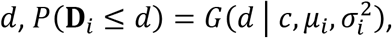, with *i*∈{A,V,AV}, depending on the modality. The expected detection time, conditional on stimulus detection before *d*, is again obtained by integration of *t.g(t)* from *t*= 0 to *d*(see Eq. 4). The solution has been originally presented in (Schwarz, 1994, Eq. 6)

In bimodal stimuli with onset asynchrony 0 < τ < d (say, without loss of generality, V(τ)A), it is necessary to distinguish the intervals [0…τ] and [*τ…d*] during which the drift (the variance) of the diffusion process amount to 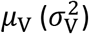 and 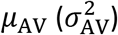, respectively. The probability for correct detections amounts to the sum of the detections within [0…τ] in which only the first stimulus contributes to the buildup of evidence; and the detections within [*τ…d*] in which both stimuli are active.

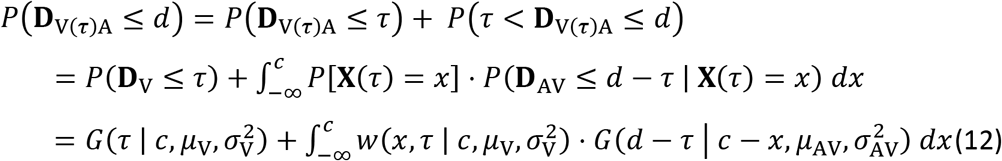

with *P[X(τ) = x] = w(x, τ*| ⋯) denoting the density of the activation of the processes not yet absorbed at *t= τ* (Eq. 5). The integrand decomposes into a sum of four terms of the form *ϕ(x)*. Φ(*x |m,s^2^*) that can be integrated using the bivariate Normal distribution (Owen, 1980, Eq. 10, 010.1, see Appendix A). For the predicted amount of correct responses, a 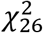 statistic is obtained (e.g.,Schwarz, 2006) by the squared difference between the predicted and observed proportion of responses, divided by the variance for binomial proportions π(1 – π)/, with π= *P*(**D** ≤ *d*) denoting the probabilities for correct detections. To avoid numerical problems close to zero or one, π was bounded within [0.01, 0.99].

The expected detection time, conditional on stimulus detection before the deadline, amounts to

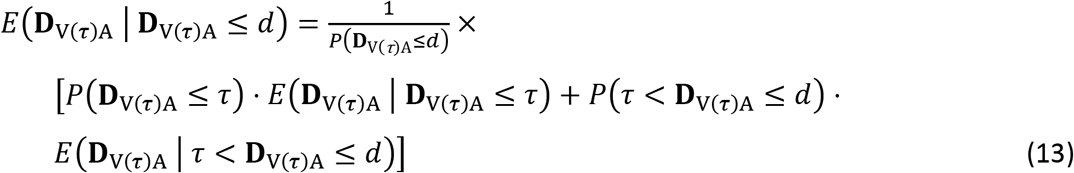

The normalization factor *P*(**D**_V*(τ)*A_ ≤ *d*) is the same as in Equation 12. The first term within the square brackets is given by Equation 4,

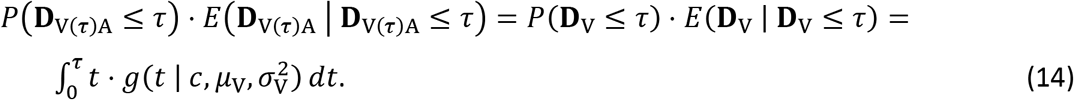

The second term is more complicated,

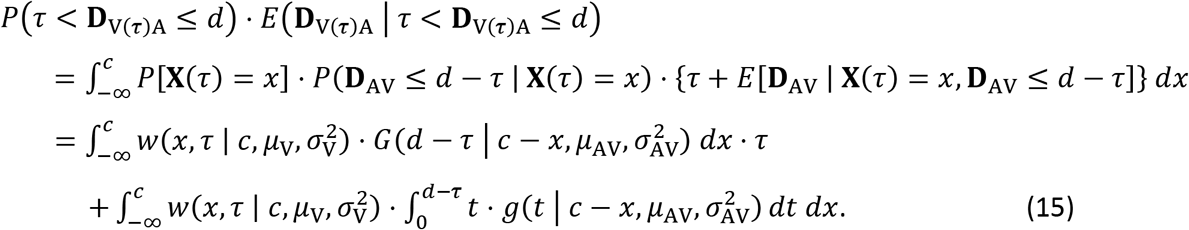

The *W·G* term corresponds to Equation 12, multiplied by τ. The double integral decomposes into four 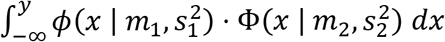 terms (Owen, 1980, Eq. 10,010.1, see Eq. 12 above) and another four terms of the form 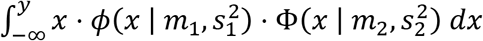 that match with (Owen, 1980, eqs. 10,010.1 and 10,011.1). Details are given in Appendix B, and R code (R Core Team, 2017) is available as online supplementary material.

For the observable mean response time, we assume again an SOA invariant mean residual μ_M_,

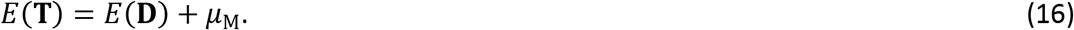

For each monkey, an approximate 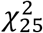 goodness-of-fit statistic can be determined by the sum of the squared standardized deviations of the predicted and the observed average response times (e.g.,Schwarz, 2006). Compared to the model without deadline, one degree of freedom is lost because the deadline is adjusted to the data. Because the *X^2^* statistics for the mean RTs and proportions of correct responses are not independent, we did not combine them but instead transformed them into *P*-values and maximized the minimum of the *P*-values as a conservative fitting criterion.

### Results for the deadline model

Figure 4 show the results from fitting the diffusion superposition model with the deadline to the behavioral performance of the monkeys (accuracy and mean RTs) as a function of the signal-to-noise ratio and the SOAs. The additional deadline parameter improves the model fits and provides a very good description of both accuracy and RTs of the monkeys. For Monkey 1 the model provided an excellent fit to the data. (Accuracy: 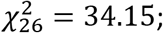 mean RT: 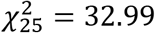, *P*_min_=0.131). In Monkey 2, the model fit was less convincing (Accuracy: 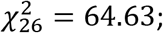 mean RT: 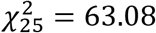, *P*_min_ < 0.001), but still acceptable given the conservative fitting procedure where we try to jointly fit both the RTs and accuracy of the monkeys. The best fit parameter estimates are shown in Table 1.

**Figure 4:**
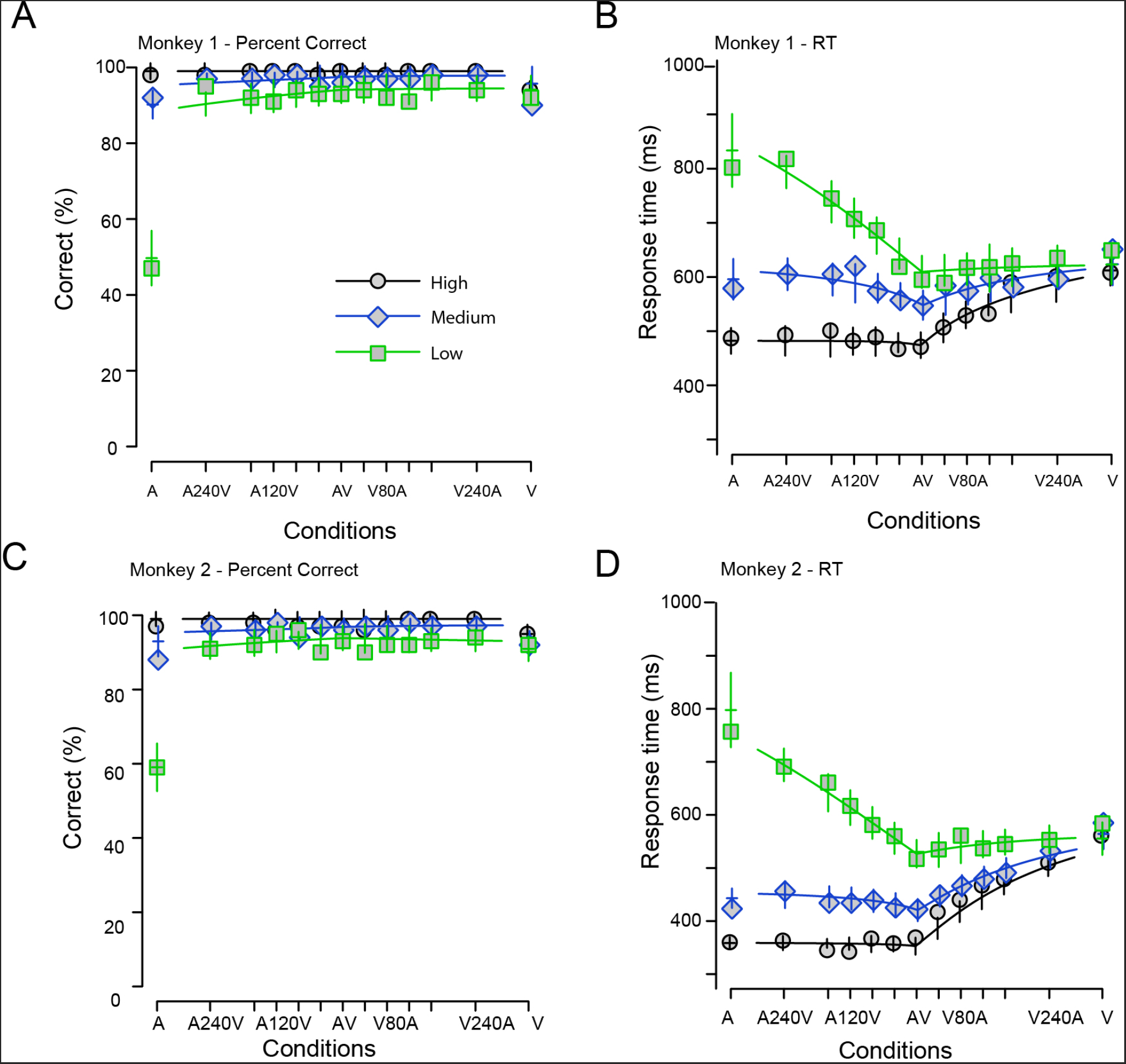
A diffusion superposition model with a time deadline predicts monkey RTs and response accuracy to audiovisual stimuli. Accuracy and RT of the two monkeys along with the fits from the diffusion superposition model with a deadline. **A, C** Accuracy of the monkeys as a function of SNR and SOA. X-axes show SOA; Y-axes show the percent correct. **B, D** Response time of the monkeys as a function of SNR And SOA. X-axes show SOA; Y-axes show Response Time in ms. In both panels, the high SNRS are shown in black circles, the medium SNRs in blue diamonds and low SNRs in green squares.

**Table 1.** Model parameters

| Parameter | Monkey 1 | Monkey 2 |
| --- | --- | --- |
| $\mu_A$ (low, medium, high) | 0.01, 0.25, 1.53 | 0.09, 0.41, 9.74 |
| $\sigma_A^2$ (low, medium, high) | 18.5, 130.3, 74.4 | 11.7, 379.6, 10.7 |
| $\mu_V$ (low, medium, high) | 0.35, 0.37, 0.34 | 0.30, 0.34, 0.37 |
| $\sigma_V^2$ (low, medium, high) | 73.2, 68.0, 92.8 | 67.8, 73.1, 41.9 |
| $\mu_M$ (msec) | 419 | 343 |
| Deadline $d$ (msec) | 951 | 828 |

## Discussion

The goal of our study was to test if the single-barrier diffusion superposition model (Schwarz, 1989, 1994; Diederich, 1995) can describe accuracy and RTs to audiovisual vocalizations of different intensities in a detection task. In the auditory modality the intensity manipulation was effective (Figures 2 and 3). In line with this, the drift estimates *μ*_A_ monotonically increased with SNR (Table 1; the variance estimates 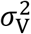 showed a less regular pattern). In the visual modality, drift estimates μ_V_ and variance estimates _V_^2^ were more or less equal for the different intensities, which is consistent with the converging pattern of the mean RTs for positive SOAs (Figure 2). The residual μ_M_ was similar in the two animals, reflecting their overall response speed and the fact that stimulus detection is probably just one of several stages of the overall response process.

Consistent with many earlier results in bimodal divided attention (Diederich & Colonius, 2004a; Molholm et al., 2002; Murray et al., 2005), separate activation (a.k.a. race) models were insufficient to explain the behavioral patterns we observed (Miller, 1982, 1986). We used many SOA conditions and thus the majority of the stimuli used in the present study were audiovisual stimuli. Enrichment of audiovisual conditions rules out trial history effects and modality shift effects as the *only* driving force of coactivation effects (Gondan, Lange, Rösler, & Röder, 2004; Miller, 1986; Otto, Dassy, & Mamassian, 2013; Otto & Mamassian, 2012).

We have focused on describing and modeling the mean RTs and mean response accuracy for a detection task across different SNRs and SOAs (Cluff et al., 2015; Crevecoeur et al., 2016; Dixon & Spitz, 1980; Holmes, 2009; Meredith et al., 1987; Stein et al., 1989; van Wassenhove et al., 2007). Some studies have addressed the effect of sensory reliability which is related to the sensory intensity manipulation we performed here on benefits of multisensory integration but did not modulate the delay between the sensory stimuli (Drugowitsch, DeAngelis, Klier, Angelaki, & Pouget, 2014). Other studies have examined the dependence on sensory delays but not on sensory intensity (Crevecoeur et al., 2016). Experiments that simultaneously vary stimulus intensity, stimulus reliability and SOA are needed to fully understand the relative roles of these factors in multisensory discrimination et al., 2010).

Our study is focused on describing the behavior of monkeys performing an audiovisual detection task. Our model is likely to apply to humans performing similar tasks. We previously showed that the behavior of monkeys and humans in a simpler version of this task that only involved variation in intensity of the sensory stimuli was very similar (Chandrasekaran et al., 2011). Our analysis here expands on a classical coactivation model which has been previously used to successfully model the behavior of a human participant across a range of SOAs (Schwarz, 1994).

We have shown that an accumulator, which integrates visual and auditory inputs to a bound, explained the behavioral benefits from multisensory integration. However, in our task design, no explicit trial onset information was provided to the animals. Instead, the stimulus arrived in a continuous ongoing stream. This paradigm has several advantages because it mimics a natural flow of stimuli in the real world and avoids sharp transients in visual stimuli. But, it raises the important question of how an integrator knows when to begin integrating the sensory evidence? One plausible solution is that a neural circuit resets the integrator after either the last behavioral action by the animal (false alarm/correct detection) or after some time has elapsed (Janssen & Shadlen, 2005). The fits may improve by incorporating previous ideas which propose to jointly model the inter stimulus interval as well as the responses to sensory stimuli.

The superposition model with a deadline predicts RTs and accuracy of monkeys when they detect *dynamic* visual and auditory stimuli (vocalizations). In other contexts, generalized variants of these coactivation models have been used with dynamic stimuli (Drugowitsch et al., 2014). We believe these types of models may also be applicable to static audiovisual stimuli for two reasons. First, Diffusion models are commonly used with static visual stimuli (Ratcliff, Thapar, & McKoon, 2003; Voss, Rothermund, & Brandtstädter, 2008). Second,the superposition model has been used to explain discrimination behavior for *static* audiovisual stimuli (Gondan et al., 2010; Schwarz, 1989, 1994). Applying these models to both static and dynamic multisensory stimuli in the same study may help test proposals that there are different mechanisms for the processing of static and dynamic multisensory stimuli (Raposo, Sheppard, Schrater, & Churchland, 2012).

The key contribution was to show that a model with additive superposition of the channel-specific evidence explains the benefits of integrating faces and voices in animal perception across a wide range of SOAs and SNRs. This class of coactivation models has previously been used to explain response times of human participants in auditory-visual detection tasks (Diederich, 1995; Schwarz, 1989, 1994). The emphasis of these additive coactivation models (or more general versions, e.g., (Drugowitsch et al., 2014)) seems *prima facie* at odds with classical reports promoting superadditive multisensory interaction (Stein & Meredith, 1993). In these studies, superadditivity, and other nonlinear mechanisms were considered fundamental for mediating benefits from multisensory integration. However, as a series of studies have shown, the majority of neurons in classical multisensory brain regions such as the superior colliculus accumulate their synaptic input in a linear manner for a range of stimulus intensities, and nonlinearities are observed only at very low intensities (Dahl, Logothetis, & Kayser, 2010; Populin & Yin, 2002; Skaliora, Doubell, Holmes, Nodal, & King, 2004; Stanford, Quessy, & Stein, 2005; Stanford & Stein, 2007; Stein & Stanford, 2008). Stated differently, additive combination is the norm. For conflicting stimuli (e.g., in temporal order judgment, where participants are asked to report which modality came first), linear summation may occur in the other direction, with the overall evidence corresponding to the difference between the channel-specific activations (Schwarz, 2006).

Besides linearity of multisensory integration in single neurons, studies increasingly demonstrate that ensembles of neurons (which might encode stimuli nonlinearly at the single neuron level) can perform linear computations (Ma, Beck, Latham, & Pouget, 2006). We believe that our abstract behavioral model might be implemented by adopting frameworks such as probabilistic population codes. For example, computationally, at the population level, linear summation of neural activation is possible and yields optimal solutions for a very general class of computational problems (Beck et al., 2008; Ma et al., 2006). Extensions of this model showed that assuming Poisson-like distributions of spike counts allows biological networks to accumulate evidence while choosing the most likely action (Beck et al., 2008). We believe our description of behavioral data by this linear summation model will assist in relating neurophysiological and modeling studies of multisensory detection and broadly integration (Chandrasekaran, 2016; Fetsch, DeAngelis, & Angelaki, 2013; Ma et al., 2006; Seilheimer et al., 2014).

## Acknowledgments

This work was supported by the National Institutes of Health (NINDS) R01NS054898 provided to Prof. Asif A. Ghazanfar (Princeton University). The experimental work was performed under his auspices in his lab. CC was supported by the Charlotte Elizabeth Procter and Centennial Fellowships from Princeton University. We thank Lauren Kelly for expert care of our monkey subjects, Shawn Steckenfinger for the creation of monkey avatars and Luis Lemus for assistance in collecting behavioral data from the monkeys. CC was supported by K99-NS092972 during the writing of this manuscript.

## Supplemental material

The online supplement to this article includes mean RTs and accuracy rates for the two animals (Tables S1–S4), bootstrap test of the race model (Tables S5–S6), as well as a commented R script with an implementation of the deadline model.

**Table S1.** Mean accuracy and bootstrap standard error for different conditions for Monkey 1

|  | A | A240V | A160V | A120V | A80V | A40V | AV | V40A | V80A | V120A | V160A | V240A | V |
| --- | --- | --- | --- | --- | --- | --- | --- | --- | --- | --- | --- | --- | --- |
| High | 98(1) | 98(1) | 99(1) | 99(1) | 99(1) | 98(1) | 99(1) | 98(1) | 98(1) | 99(1) | 99(1) | 99(1) | 94(2) |
| Medium | 92(2) | 97(1) | 97(1) | 98(1) | 98(1) | 95(2) | 96(1) | 97(1) | 97(1) | 97(1) | 98(1) | 98(1) | 90(2) |
| Low | 47(4) | 95(2) | 92(2) | 91(2) | 94(2) | 93(2) | 93(2) | 94(2) | 92(2) | 91(2) | 96(2) | 94(2) | 92(2) |

**Table S2.** Mean RT and Standard Error of RTs for different conditions for Monkey 1.

|  | A | A240V | A160V | A120V | A80V | A40V | AV | V40A | V80A | V120A | V160A | V240A | V |
| --- | --- | --- | --- | --- | --- | --- | --- | --- | --- | --- | --- | --- | --- |
| High | 487(12) | 493(14) | 501(15) | 482(13) | 489(14) | 467(9) | 471(12) | 507(14) | 529(13) | 532(10) | 590(15) | 601(17) | 608(14) |
| Medium | 579(19) | 605(15) | 605(15) | 620(18) | 574(14) | 557(12) | 547(14) | 584(18) | 574(15) | 598(17) | 581(14) | 596(14) | 651(20) |
| Low | 802(34) | 818(15) | 745(20) | 707(19) | 686(17) | 619(15) | 595(15) | 589(15) | 617(15) | 618(22) | 625(18) | 635(20) | 649(19) |

**Table S3.**
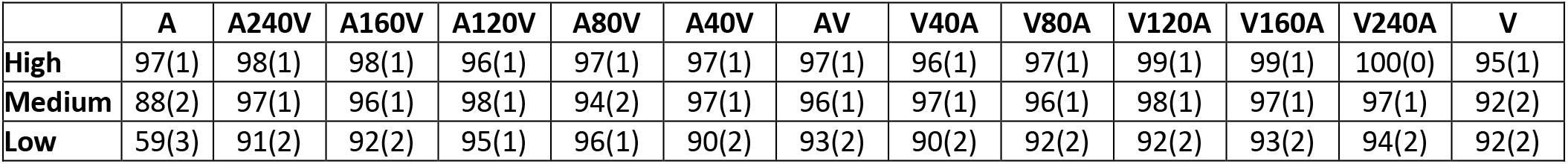
Mean accuracy and bootstrap standard error for different conditions for Monkey 2

**Table S4.**
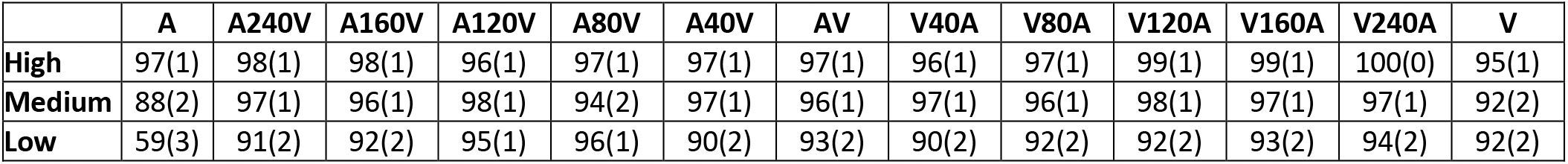
Mean RT and Standard Error of RTs for different conditions for Monkey 2.

**Table S5.** *P* values for race model test (Monkey 1)

|  | A240V | A160V | A120V | A80V | A40V | AV | V40A | V80A | V120A | V160A | V240A |
| --- | --- | --- | --- | --- | --- | --- | --- | --- | --- | --- | --- |
| High | 0.057 | 0.062 | 0.014 | 0.039 | 0.038 | 0.031 | 0.018 | 0.000 | 0.000 | 0.000 | 0.000 |
| Medium | 0.125 | 0.013 | 0.029 | 0.062 | 0.293 | 0.002 | 0.000 | 0.000 | 0.000 | 0.000 | 0.000 |
| Low | 0.043 | 0.000 | 0.000 | 0.000 | 0.000 | 0.001 | 0.000 | 0.040 | 0.000 | 0.007 | 0.000 |

**Table S6.** *P* values for race model test for Monkey 2

|  | A240V | A160V | A120V | A80V | A40V | AV | V40A | V80A | V120A | V160A | V240A |
| --- | --- | --- | --- | --- | --- | --- | --- | --- | --- | --- | --- |
| High | 0.001 | 0.001 | 0.000 | 0.004 | 0.005 | 0.033 | 0.000 | 0.000 | 0.000 | 0.000 | 0.000 |
| Medium | 0.008 | 0.003 | 0.011 | 0.041 | 0.000 | 0.000 | 0.004 | 0.000 | 0.000 | 0.000 | 0.000 |
| Low | 0.019 | 0.001 | 0.000 | 0.000 | 0.000 | 0.000 | 0.000 | 0.000 | 0.000 | 0.001 | 0.000 |

## Appendix A: Probability of absorption

Here we derive the explicit expressions for the probability of absorption in bimodal stimuli with onset asynchrony 0 < τ < d(Eq. 12). Without loss of generality, we consider the case V(τ)A in which the visual stimulus is presented first. Between *t* = 0 and *t* = τ, only the visual channel contributes to the build-up of evidence, so the probability of absorption within the interval (0, τ) is given by

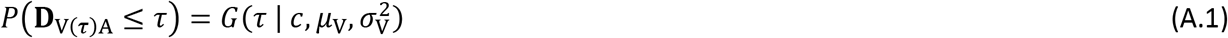

with *G* denoting the inverse Gaussian distribution (Eq. 2). Later, within the time interval (*τ,d*), the probability of absorption is the mixture of absorption probabilities of those processes still active at time τ, with the barrier depending on the activation **X**(τ) <*c*, weighted by the density of processes at **X**(τ). Let *d’ =d –τ*. Then,

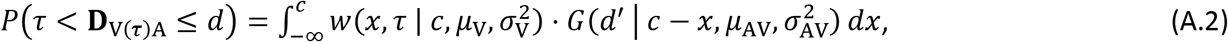

with *w* given by Equation 5. The integrand in (A.2) can be transformed into four integrals of the form 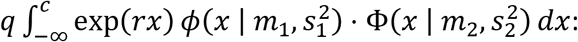

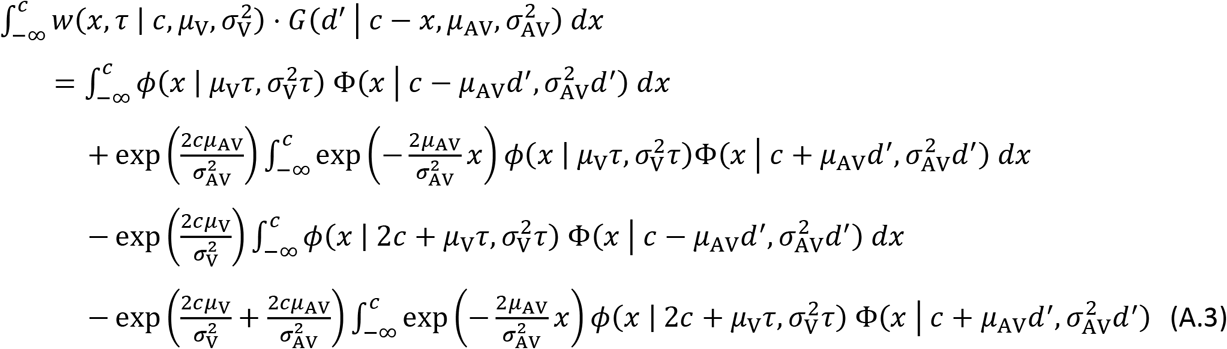

By completing the square, we have exp 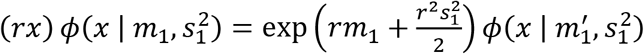, with 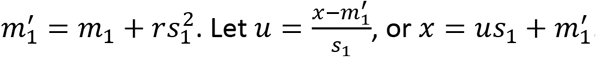 then, the integral can be rewritten as

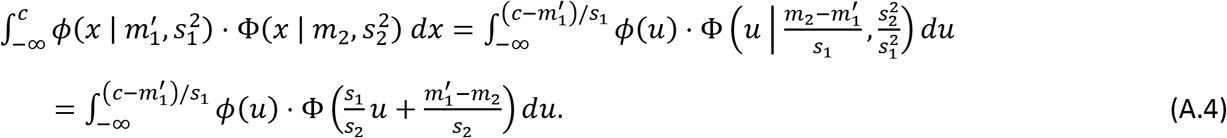

The form of (A.4) now matches Owen (1980, Eq. 10,010.1) which can be determined by the bivariate Normal distribution, 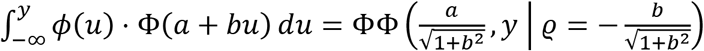. An implementation in R (R Core Team, 2017) is available as online supplementary material.

## Appendix B: Conditional mean response time

Here we derive the explicit expressions for the conditional mean response time for bimodal stimuli with onset asynchrony τ, conditional on absorption before the deadline (Eqs. 13–15). We consider again the case V(τ)A in which the visual stimulus is presented first. Between *t* = 0 and *t= τ*, only the visual channel contributes to the build-up of evidence, so the conditional mean RT is given by Schwarz (1994, Equation A.2).

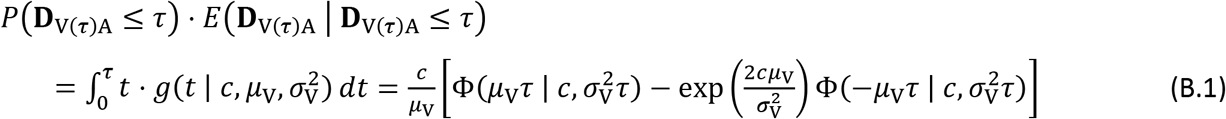

Later, within the time interval (*τ,d*), the expected detection time is again a mixture of expected detection times for the processes still active at time τ, weighted by the density of processes at **X**(τ). These processes now have increased drift μ_AV_ and variance 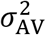 the work to be done (i.e., the barrier) depends on the activation **X**(τ) <*c*. Note that τ milliseconds have already passed since stimulus onset, hence the remaining time is *d^′^ = d-τ*.

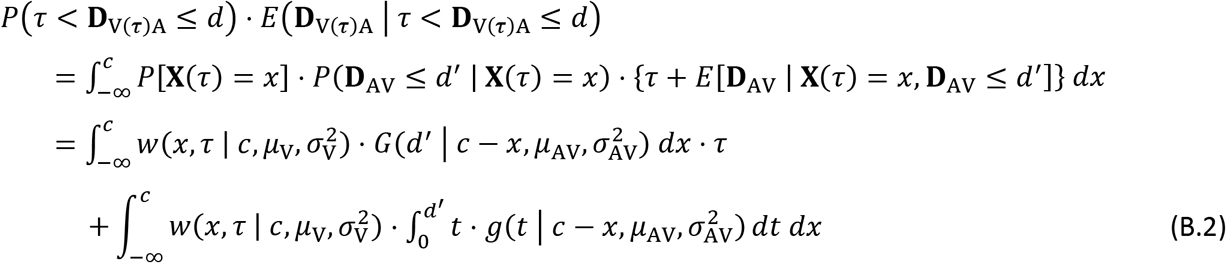

The first term corresponds exactly to (A.2), multiplied by the onset asynchrony τ. See again Schwarz (1994, Equation A.2) for 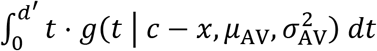 the double integral in (B.2) can then be rewritten as

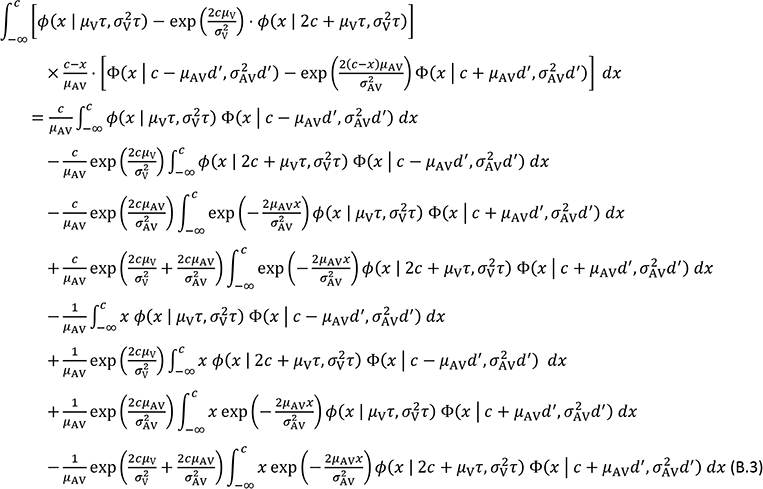

The first four terms correspond to (A.3), multiplied by a constant (± *c*/μ_AV_). By completing the square, we have again exp 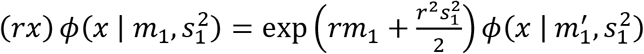 with 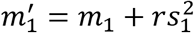 Let 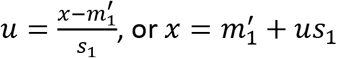 Then, the integral can be rewritten as

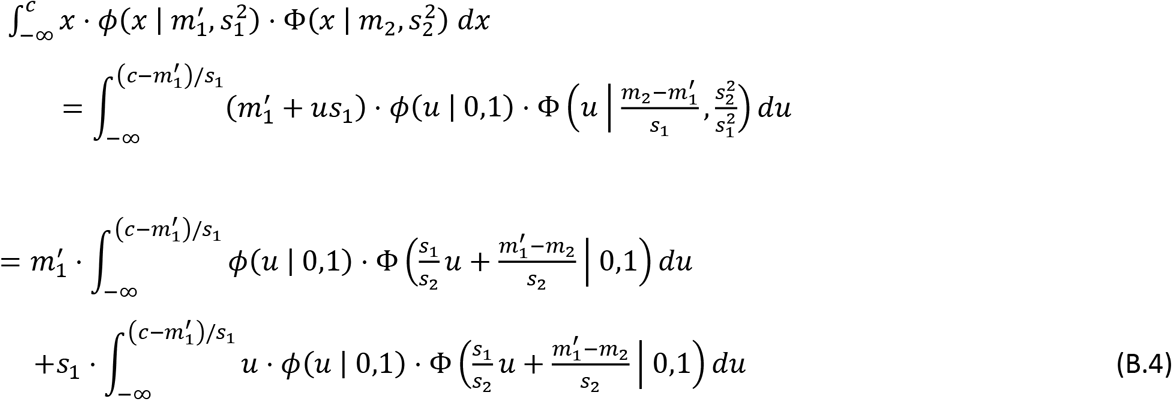

The first term of (B.4) matches again (Owen, 1980, Eq. 10,010.1). The second term matches Eq. 10,010.1 from (Owen, 1980, Eq. 10,010.1) and is calculated by

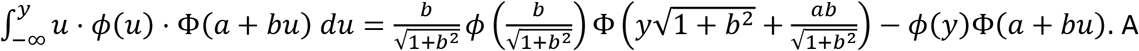

implementation in R (R Core Team, 2017) is available as an online supplement.

## Author Contributions

CC and MG designed the experiment. CC performed experiments. CC and MG analyzed the data. MG developed the formulation of the deadline model. CC and MG wrote the paper. Both authors evaluated data and modeling results.

**Accuracy and mean response time in a diffusion superposition model with deadline**

*Online supplement for “Audiovisual detection at different intensities and delays”*

This online supplement provides implementation details on the derivation of the predictions for mean response time and accuracy for the diffusion superposition model (Diederich, 1995; Schwarz, 1994) with a deadline. For simplicity, we reiterate the relevant parts of the methods section here and then add code in R statistical language for the different equations. The R code (R Core Team, 2017) includes the necessary defaults that allow testing and deployment in other analyses.

## Libraries

The code requires the inverse Gaussian distribution package SuppDists (Wheeler, 2013, available from CRAN). In addition, package mvtnorm (Genz et al., 2014) is used for the bivariate Normal distribution.

~~~
#
~~~

~~~
# Implementation of the inverse Gaussian distribution (Wheeler, 2016, install from CRAN first)
~~~

~~~
#
~~~

~~~
library(SuppDists)
~~~

~~~
#
~~~

~~~
# Multivariate Normal distribution (Genz et al., 2016)
~~~

~~~
# library(mvtnorm)
~~~

## Accuracy

For unimodal/synchronous stimuli, accuracy is given by the inverse Gaussian distribution at the deadline *d*. For example, Monkey 1’s accuracy in Condition v (low intensity) is given by acc_sync(d=1000, c=100, mu=0.13, sigma2=4.3^2).

~~~
#
~~~

~~~
# Accuracy in unimodal and synchronous stimuli
~~~

~~~
#
~~~

~~~
acc_sync = function(d, c, mu, sigma2)
~~~

~~~
{
~~~

~~~
pinvGauss(d, nu=c/mu, lambda=c*c/sigma2)
~~~

~~~
}
~~~

For stimuli with onset asynchrony τ, accuracy is given by (12) which is the sum of the inverse Gaussian distribution at time τ and four integrals of the form 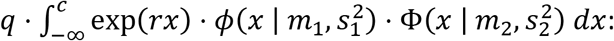

1. 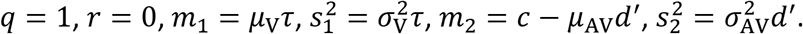
2. 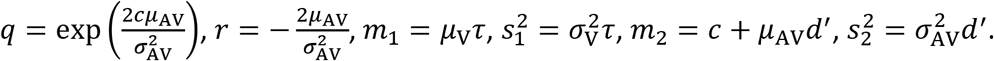
3. 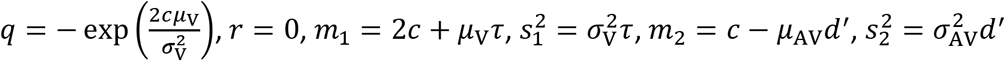
4. 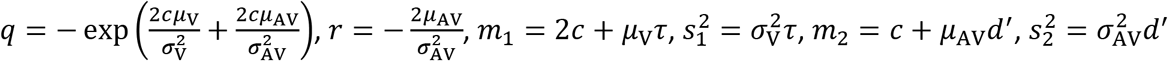

with *d’ = d – τ* Three convenience functions evaluate these integrals using Eq. 10,010.1 in (Owen, 1980):

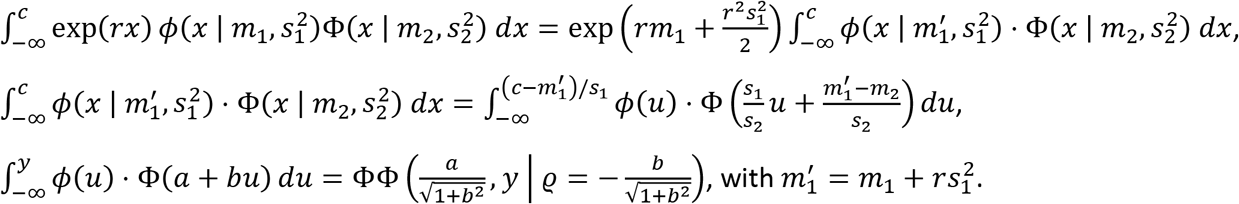

~~~
#
~~~

~~~
# Integrate dnorm(x) * pnorm(a + b*x) from-Inf to y (Owen, 1980, Eq. 10,010.1)
~~~

~~~
#
~~~

~~~
owen10_010.1 = function(y, a, b)
~~~

~~~
{
~~~

~~~
rho =-b/sqrt(1 + b*b)
~~~

~~~
pmvnorm(lower=c(-Inf,-Inf), upper=c(a/sqrt(1+b*b), y), corr=matrix(c(1, rho, rho, 1), nrow=2))
~~~

~~~
}
~~~

~~~
#
~~~

~~~
# Integrate dnorm(x | mu1, sigma1) * pnorm(x | mu2, sigma2) from-Inf to y
~~~

~~~
#
~~~

~~~
owen10_010.1b = function(y, mu1, sigma1, mu2, sigma2)
~~~

~~~
{
~~~

~~~
owen10_010.1(y=(y – mu1)/sigma1, a=(mu1 – mu2)/sigma2, b=sigma1/sigma2)
~~~

~~~
}
~~~

~~~
#
~~~

~~~
# Integrate exp(rx) * dnorm(x | mu1, sigma1) * pnorm(x | mu2, sigma2) from-Inf to y
~~~

~~~
#
~~~

~~~
owen10_010.1c = function(y, r, mu1, sigma1, mu2, sigma2)
~~~

~~~
{
~~~

~~~
exp(r * mu1 + r^2 * sigma1^2 / 2) * owen10_010.1b(y, mu1 + r * sigma1^2, sigma1, mu2, sigma2)
~~~

~~~
}
~~~

~~~
acc_async = function(d, c, mua, sigma2a, mub, sigma2b, tau)
~~~

~~~
{
~~~

~~~
muab = mua + mub
~~~

~~~
sigma2ab = sigma2a + sigma2b
~~~

~~~
# Probability of absorption within 0…tau
~~~

~~~
p0 = acc_sync(d=tau, c=c, mu=mua, sigma2=sigma2a)
~~~

~~~
# 1st term of Equation A.3
~~~

~~~
q = 1
~~~

~~~
r = 0
~~~

~~~
mu1 = mua * tau
~~~

~~~
sigma1=sqrt(sigma2a * tau)
~~~

~~~
mu2 = c – muab * (d – tau)
~~~

~~~
sigma2 = sqrt(sigma2ab * (d – tau))
~~~

~~~
p1 = q * owen10_010.1c(c, r, mu1, sigma1, mu2, sigma2)
~~~

~~~
# 2nd integral
~~~

~~~
q = exp(2 * c * muab / sigma2ab)
~~~

~~~
r =-2 * muab / sigma2ab
~~~

~~~
mu1 = mua * tau
~~~

~~~
sigma1 = sqrt(sigma2a * tau)
~~~

~~~
mu2 = c + muab * (d – tau)
~~~

~~~
sigma2 = sqrt(sigma2ab * (d – tau))
~~~

~~~
p2 = q * owen10_010.1c(c, r, mu1, sigma1, mu2, sigma2)
~~~

~~~
# 3rd integral
~~~

~~~
q =-exp(2 * c * mua / sigma2a)
~~~

~~~
r = 0
~~~

~~~
mu1 = 2 * c + mua * tau
~~~

~~~
sigma1 = sqrt(sigma2a * tau)
~~~

~~~
mu2 = c – muab * (d – tau)
~~~

~~~
sigma2 = sqrt(sigma2ab * (d – tau))
~~~

~~~
p3 = q * owen10_010.1c(c, r, mu1, sigma1, mu2, sigma2)
~~~

~~~
# 4th integral
~~~

~~~
q =-exp(2 * c * mua / sigma2a + 2 * c * muab / sigma2ab)
~~~

~~~
r =-2 * muab / sigma2ab
~~~

~~~
mu1 = 2 * c + mua * tau
~~~

~~~
sigma1 = sqrt(sigma2a * tau)
~~~

~~~
mu2 = c + muab * (d – tau)
~~~

~~~
sigma2 = sqrt(sigma2ab * (d – tau))
~~~

~~~
p4 = q * owen10_010.1c(c, r, mu1, sigma1, mu2, sigma2)
~~~

~~~
p0 + p1 + p2 + p3 + p4
~~~

~~~
}
~~~

For example, Monkey 1’s accuracy in Condition v100a (low intensity) is given by acc_async(d=1000, c=100, mua=0.13, sigma2a=4.3^2, mub=0.34, sigma2b=11.7^2, tau=67).

## Mean response time

For unimodal/synchronous stimuli, the mean RT is given by Equation 4 which integrates the product of the time and the inverse Gaussian density from zero until the deadline *d*. For example, Monkey 1’s mean RT in Condition v (low intensity) is given by mrt_sync(d=1000, c=100, mu=0.13, sigma2=4.3^2).

~~~
#
~~~

~~~
# Mean RT in unimodal and synchronous stimuli
~~~

~~~
#
~~~

~~~
mrt_sync = function(d, c, mu, sigma2)
~~~

~~~
{
~~~

~~~
# Integral t * density from 0 to d (Schwarz, 1994, Equation A.2)
~~~

~~~
m = c / mu * {pnorm(mu*d, c, sqrt(sigma2*d))-
~~~

~~~
exp(2*c*mu / sigma2) * pnorm(pnorm(-mu*d, c, sqrt(sigma2*d))}
~~~

~~~
# Normalize with detection accuracy
~~~

~~~
m / acc_sync(d, c, mu, sigma2)
~~~

~~~
}
~~~

For stimuli with onset asynchrony τ, mean RT is given by Equations 13–15 which boil down to Equations 4 and 12, and four integrals of the form 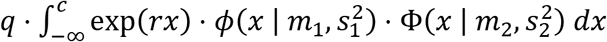, as well as four integrals of the form 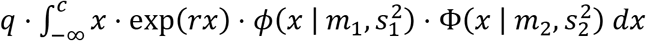

1. 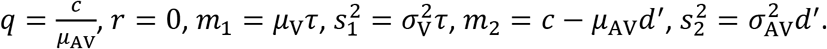
2. 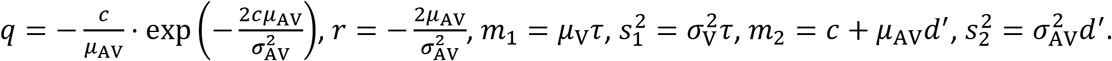
3. 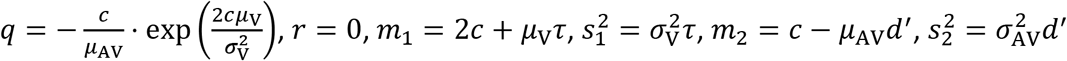
4. 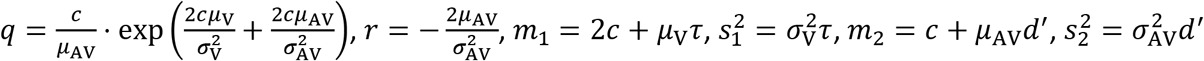
5. 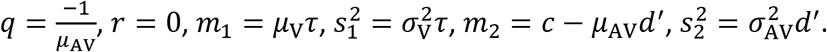
6. 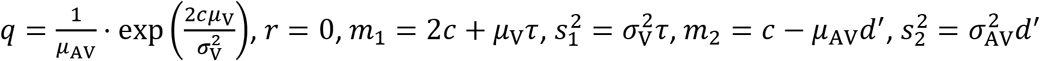
7. 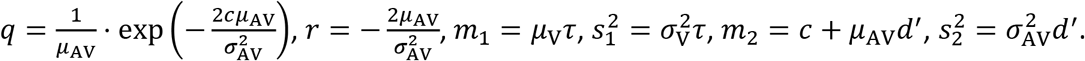
8. 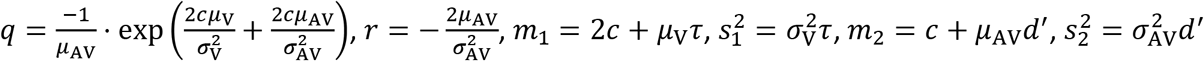

We defined again convenience functions that transform these integrals to two expressions that match Equations 10,010.1 and 10,011.1 in (Owen, 1980):

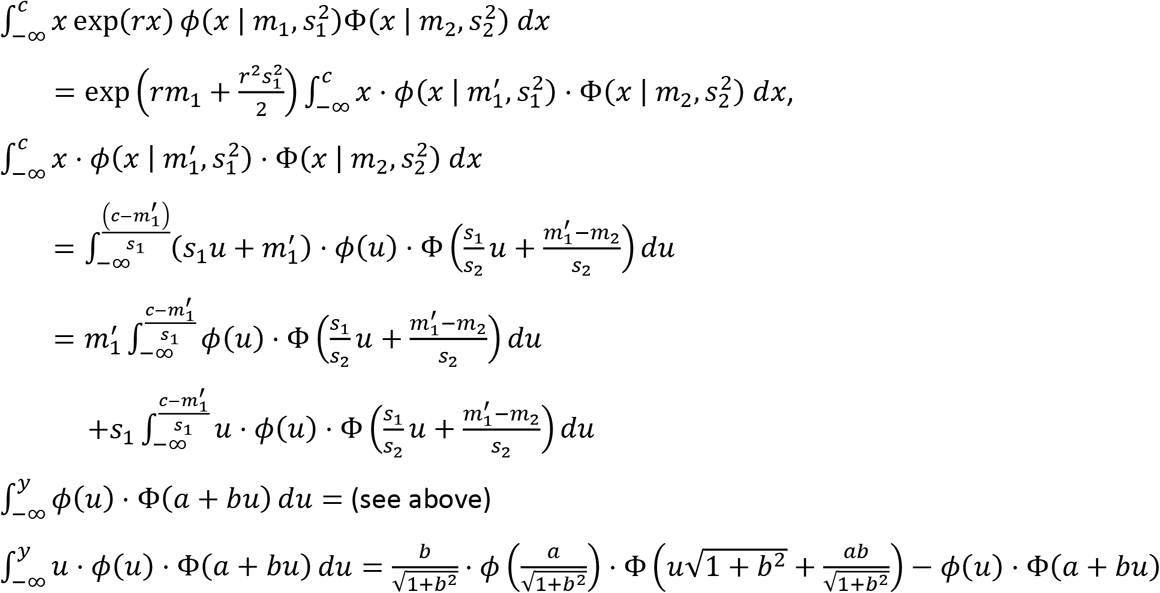

~~~
#
~~~

~~~
# Integrate dnorm(x) * pnorm(a + b*x) from-Inf to y (Owen, 1980, Eq. 10,011.1)
~~~

~~~
#
~~~

~~~
owen10_011.1 = function(y, a, b)
~~~

~~~
{
~~~

~~~
bb = sqrt(1 + b*b)
~~~

~~~
b/bb * dnorm(a/bb) * pnorm(y*bb + a*b/bb) – pnorm(a + b*y)*dnorm(y)
~~~

~~~
}
~~~

~~~
#
~~~

~~~
# Integrate x * dnorm(x | mu1, sigma1) * pnorm(x | mu2, sigma2) from-Inf to y
~~~

~~~
#
~~~

~~~
owen10_011.1b = function(y, mu1, sigma1, mu2, sigma2)
~~~

~~~
{
~~~

~~~
owen10_011.1(y=(y – mu1)/sigma1, a=(mu1 – mu2)/sigma2, b=sigma1/sigma2) * sigma1 +
~~~

~~~
mu1 * owen10_010.1(y=(y – mu1)/sigma1, a=(mu1 – mu2)/sigma2, b=sigma1/sigma2)
~~~

~~~
}
~~~

~~~
#
~~~

~~~
# Integrate x * exp(rx) * dnorm(x | mu1, sigma1) * pnorm(x | mu2, sigma2) from-Inf to y
~~~

~~~
#
~~~

~~~
owen10_010.1c = function(y, r, mu1, sigma1, mu2, sigma2)
~~~

~~~
{
~~~

~~~
exp(mu1 * v + sigma1^2 * v^2 / 2) * owen10_011.1b(y, mu1 + v * sigma1^2, sigma1, mu2, sigma2)
~~~

~~~
}
~~~

For example, Monkey 1’s mean RT in Condition v100a (low intensity) is predicted to mrt_async(d=1000, c=100, mua=0.13, sigma2a=4.3^2, mub=0.34, sigma2b=11.7^2, tau=67).

~~~
mrt_async = function(d, c, mua, sigma2a, mub=0.34, sigma2b, tau)
~~~

~~~
{
~~~

~~~
muab = mua + mub
~~~

~~~
sigma2ab = sigma2a + sigma2b
~~~

~~~
d_ = d – tau
~~~

~~~
# Integral t * density from 0 to tau (Schwarz, 1994, Equation A.2)
~~~

~~~
m0 = c / mua * {pnorm(mua*tau, c, sqrt(sigma2a*tau))-
~~~

~~~
exp(2*c*mua / sigma2a) * pnorm(pnorm(-mua*tau, c, sqrt(sigma2a*tau))}
~~~

~~~
# First term of Eq. 15 (tau * second term of Eq. 12)
~~~

~~~
mtau = tau*{acc_async(d, c, mua, sigma2a, mub, sigma2b, tau) – acc_sync(d, c, mua, sigma2a)}
~~~

~~~
# 1st integral in B.3
~~~

~~~
q = c / mua
~~~

~~~
r = 0
~~~

~~~
mu1 = mua * tau
~~~

~~~
sigma1=sqrt(sigma2a * tau)
~~~

~~~
mu2 = c – muab * d_
~~~

~~~
sigma2 = sqrt(sigma2ab * d_)
~~~

~~~
m1 = q * owen10_010.1c(c, r, mu1, sigma1, mu2, sigma2)
~~~

~~~
# 2nd integral
~~~

~~~
q =-c / muab * exp(2 * c * muab / sigma2ab)
~~~

~~~
r =-2 * muab / sigma2ab
~~~

~~~
mu1 = mua * tau
~~~

~~~
sigma1 = sqrt(sigma2a * tau)
~~~

~~~
mu2 = c + muab * d_
~~~

~~~
sigma2 = sqrt(sigma2ab * d_)
~~~

~~~
m2 = q * owen10_010.1c(c, r, mu1, sigma1, mu2, sigma2)
~~~

~~~
# 3rd integral
~~~

~~~
q =-c / muab * exp(2 * c * mua / sigma2a)
~~~

~~~
r = 0
~~~

~~~
mu1 = 2 * c + mua * tau
~~~

~~~
sigma1 = sqrt(sigma2a * tau)
~~~

~~~
mu2 = c – muab * d_
~~~

~~~
sigma2 = sqrt(sigma2ab * d_)
~~~

~~~
m3 = q * owen10_010.1c(c, r, mu1, sigma1, mu2, sigma2)
~~~

~~~
# 4th integral
~~~

~~~
q = c / muab * exp(2 * c * mua / sigma2a + 2 * c * muab / sigma2ab)
~~~

~~~
r =-2 * muab / sigma2ab
~~~

~~~
mu1 = 2 * c + mua * tau
~~~

~~~
sigma1 = sqrt(sigma2a * tau)
~~~

~~~
mu2 = c + muab * d_
~~~

~~~
sigma2 = sqrt(sigma2ab * d_)
~~~

~~~
m4 = q * owen10_010.1c(c, r, mu1, sigma1, mu2, sigma2)
~~~

~~~
# 5th integral in B.3
~~~

~~~
q =-1 / mua
~~~

~~~
r = 0
~~~

~~~
mu1 = mua * tau
~~~

~~~
sigma1 = sqrt(sigma2a * tau)
~~~

~~~
mu2 = c – muab * d_
~~~

~~~
sigma2 = sqrt(sigma2ab * d_)
~~~

~~~
m5 = q * owen10_011.1c(c, r, mu1, sigma1, mu2, sigma2)
~~~

~~~
# 6th integral
~~~

~~~
q = 1 / muab * exp(2 * c * muab / sigma2ab)
~~~

~~~
r =-2 * muab / sigma2ab
~~~

~~~
mu1 = mua * tau
~~~

~~~
sigma1 = sqrt(sigma2a * tau)
~~~

~~~
mu2 = c + muab * d_
~~~

~~~
sigma2 = sqrt(sigma2ab * d_)
~~~

~~~
m6 = q * owen10_011.1c(c, r, mu1, sigma1, mu2, sigma2)
~~~

~~~
# 7th integral
~~~

~~~
q = 1 / muab * exp(2 * c * mua / sigma2a)
~~~

~~~
r = 0
~~~

~~~
mu1 = 2 * c + mua * tau
~~~

~~~
sigma1 = sqrt(sigma2a * tau)
~~~

~~~
mu2 = c – muab * d_
~~~

~~~
sigma2 = sqrt(sigma2ab * d_)
~~~

~~~
m7 = q * owen10_011.1c(c, r, mu1, sigma1, mu2, sigma2)
~~~

~~~
# 8th integral
~~~

~~~
q =-1 / muab * exp(2 * c * mua / sigma2a + 2 * c * muab / sigma2ab)
~~~

~~~
r =-2 * muab / sigma2ab
~~~

~~~
mu1 = 2 * c + mua * tau
~~~

~~~
sigma1 = sqrt(sigma2a * tau)
~~~

~~~
mu2 = c + muab * d_
~~~

~~~
sigma2 = sqrt(sigma2ab * d_)
~~~

~~~
m8 = q * owen10_011.1c(c, r, mu1, sigma1, mu2, sigma2)
~~~

~~~
# Return value: normalized integral t * density from 0 to d (Eq. 13)
~~~

~~~
p = acc_async(d, c, mua, sigma2a, mub, sigma2b, tau)
~~~

~~~
(m0 + mtau + m1 + m2 + m3 + m4 + m5 + m6 + m7 + m8) / p
~~~

~~~
}
~~~

